# Bayes-Like Integration of a New Sensory Skill with Vision

**DOI:** 10.1101/232579

**Authors:** James Negen, Lisa Wen, Lore Thaler, Marko Nardini

## Abstract

Humans are effective at dealing with noisy, probabilistic information in familiar settings. One hallmark of this is Bayesian Cue Combination: combining multiple noisy estimates to increase precision beyond the best single estimate, taking into account their reliabilities. Here we show that adults also combine a novel audio cue to distance, akin to human echolocation, with a visual cue. Following two hours of training, subjects were more precise given both cues together versus the best single cue. This persisted when we changed the novel cue’s auditory frequency. Reliability changes also led to a re-weighting of cues without feedback, showing that they learned something more flexible than a rote decision rule for specific stimuli. The main findings replicated with a vibrotactile cue. These results show that the mature sensory apparatus can learn to flexibly integrate new sensory skills. The findings are unexpected considering previous empirical results and current models of multisensory learning.

## Bayes-Like Integration of a New Sensory Skill with Vision

Using the information from our senses to make the best decisions can be surprisingly complex. Consider the simple act of comparing the distances of two free supermarket checkouts. There are potentially a dozen different methods of estimating visual distance (Howard & Rogers, 2008), such as using perspective information, using the differences in the two eyes’ views of the same scene, comparing the apparent sizes of counters and other nearby objects whose sizes are familiar, and so on. From a dozen different methods, we might get a dozen different estimates. Since we can only act out one decision, one singular estimate must be decided upon. To arrive at a single estimate, different techniques can be pursued, some of which might be more prone to fail. For example, picking one estimation method at random could be extremely inaccurate if an unreliable method were selected. Averaging all estimates together might also result in a poor decision if a few very bad estimates and a few very good estimates were given equal weight. Even if we somehow knew that one estimation method is most reliable and decided to use it just by itself, we would be throwing away a lot of potentially useful information. How do we best handle all of this sensory input?

One good solution is to consider not just the individual estimates, but also their error distributions (Ernst & Banks, 2002). If the different methods give approximately normally-distributed (Gaussian) error, the process for forming a single unified estimate with the lowest variance (uncertainty) is to take an average that is weighted by each estimate’s precision (1/variance). This strategy is used explicitly by statisticians for meta-analysis and by engineers developing sensing and control systems (Martin, Fowlkes, & Malik, 2004). Surprisingly, human adults’ behaviour suggests that their perceptual systems also closely approximate this type of computation when making perceptual decisions using multiple familiar sensory inputs (Alais & Burr, 2004; Ernst & Banks, 2002; Hillis et al., 2004; Knill & Saunders, 2003). To achieve this, perceptual systems must represent the reliabilities of their estimates. The algorithm for reliability-weighted averaging can be broadly termed *Bayes-Like Cue Combination* since it is considered optimal under Bayesian statistics, and also because it is widely associated with modelling perception as approximating Bayesian inference, which provides a coherent explanation for diverse findings in the study of perception and cognition (Knill & Pouget, 2004; Pouget, Beck, Ma, & Latham, 2013).

Here we ask whether people’s previously-documented Bayes-like cue combination abilities extend beyond the use of highly familiar sensory cues, such as visual and haptic cues to object size (Ernst & Banks, 2002), into the realm of sensory substitution and augmentation, where people are trained to use new techniques or devices to perceive their environment (Maidenbaum & Abboud, 2014; Thaler & Goodale, 2016). Understanding adults’ capacities for integrating newly-learned signals with existing ones has important applications to expanding the human sensory repertoire. What if an augmented sense could be combined with existing senses, so enhancing existing sensory skills instead of replacing them? If so, people with moderate vision loss could not only be taught echolocation (Kolarik, Cirstea, Pardhan, & Moore, 2014; Stroffregen & Pittenger, 1995; Thaler & Goodale, 2016) or use of a sensory substitution device (Abboud, Hanassy, Levy-Tzedek, Maidenbaum, & Amedi, 2014; Maidenbaum et al., 2014; Meijer, 1992), but also have it combine with remaining vision, gaining something that is better than either the augmentation or the remaining vision alone. A surgeon could learn new, auditory cues to their instrument’s position or the material it is contacting, combining the audio and visual information to perform procedures more accurately. Applications such as these depend not just on learning an augmented sensory skill, which we already know people can do, but also combining it with pre-existing sensory skills in an efficient Bayes-like manner.

Can people actually do this? We are not aware of any empirical demonstration that they can. In addition, current theoretical models of multisensory perceptual learning suggest that learning to combine an augmented sense would be a truly extraordinary effort – essentially impossible given the limited resources of a routine laboratory experiment or a routine therapy method. Specifically, they suggest that it would take a full decade of daily cue-specific experience for Bayes-like combination to emerge (Daee, Mirian, Ahmadabadi, Brenner, & Tenenbaum, 2014; Weisswange, Rothkopf, Rodemann, & Triesch, 2011). This view may explain why children under 10-12 years don’t yet combine sensory cues in this way (Gori, Del Viva, Sandini, & Burr, 2008; Gori, Sandini, & Burr, 2012; Nardini, Jones, Bedford, & Braddick, 2008; Petrini, Remark, Smith, & Nardini, 2014), even within the same modality (Dekker et al., 2015; Nardini, Bedford, & Mareschal, 2010), despite having a number of other skills in the domain of cross-modal sensory perception (Bahrick & Lickliter, 2000; Gottfried, Rose, & Bridger, 1977; Lewkowicz, 2000; Lewkowicz & Turkewitz, 1980; Spelke, 1979).

On the other hand, there is evidence that probabilistic sensory-motor computations (Knill & Pouget, 2004; Pouget et al., 2013) can adapt flexibly. For example, participants quickly learn new prior distributions and adapt their weighting of these distributions in a Bayes-like manner (Bejjanki, Knill, & Aslin, 2016; Körding & Wolpert, 2004). Thus, there might perhaps be more flexibility than suggested by the models reviewed above (Daee et al., 2014; Weisswange et al., 2011). In other words, an alternative hypothesis is that humans can actually learn to combine novel sensory cues, in line with the other kinds of adaptation that have already been documented (Knill & Pouget, 2004; Pouget et al., 2013).

To examine this empirically, we developed a new experimental task to train people to use a new sensory cue to distance (Figure 1). Adult participants, wearing headphones and head-mounted displays, were immersed in a virtual environment. They learned to estimate how far away a virtual cartoon whale was hiding. Our new cue to distance was inspired by human echolocation, a technique of listening to reflected sound to perceive surrounding objects and their spatial layout (Kolarik et al., 2014; Stroffregen & Pittenger, 1995; Thaler & Goodale, 2016). The full echo signal was reduced to the echo delay component: a longer time between the initial sound and its “echo” means that the target is farther away. The delay was based on the actual speed of sound in air, approximated at 350m/s. We also showed participants how to use an independent noisy visual cue, a display of ‘bubbles’ in which a wider point in the display meant that the whale was more likely to be there. The crucial tests then examined how participants performed when both cues were presented simultaneously.

**Figure 1.**
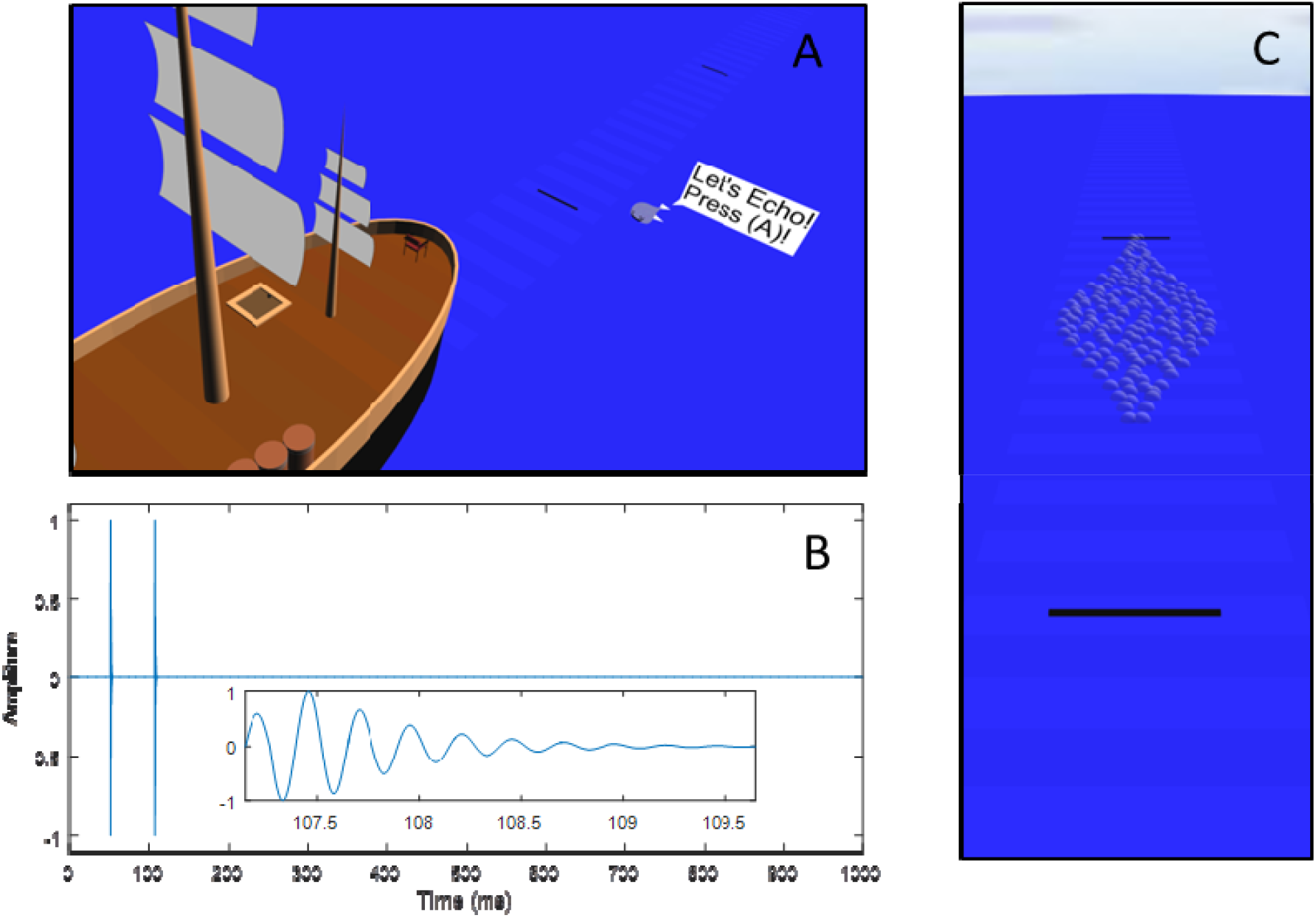
(A): The virtual environment (note that in the actual experiment participants saw the environment stereoscopically via a headset). Participants were asked to indicate how far along the line the whale was hiding based on audio and/or visual cues. (B): An example audio stimulus, with a zoom image of 2.5ms of the echo inlaid. In this case, the onset delay is about 57ms so the whale is about 10m away. (C): An example of the noisy visual stimulus, a display of bubbles. The whale is more likely to be at distances at which the patch is wider (has more bubbles). Given this cue alone, the best strategy is just to respond where it is widest. Given both cues together, the optimal strategy involves weighting this point of highest density with the location perceived through the echo-like cue.

Over a series of five sessions lasting about one hour each, we first trained distance perception via the novel cue, then tested for two central markers of Bayes-like combination with vision. The first criterion is reducing uncertainty (variable error) below the best single estimate when both cues are available (Ernst & Banks, 2002). The second criterion is flexibly re-weighting (changing reliance on) different cues when they change in reliability, even without feedback on trials with both cues (Maloney & Mamassian, 2009). The presence of both these markers together suggests that the two cues are being averaged in a way that accounts for their reliabilities, gaining precision over the best possible use of either single cue alone, and rules out alternative models based on rote learning or weaker cue interactions. If naïve adults show both markers, we will have evidence for the acquisition of Bayes-like cue combination with an augmented sense. This would lead to revising current models of multisensory learning (Daee et al., 2014; Weisswange et al., 2011) and would have important potential applications to augmenting the human senses.

## Results

### Summary

In short, we found consistent positive results in favour of Bayes-like cue combination. The five sessions were structured as follows: healthy adult participants were first asked to learn how to use an echo-like cue to distance (Sessions 1 and 2); then to also use a noisy visual cue to distance (Session 2); then to also use both simultaneously (Session 3); and finally to generalize their learning to a new audio frequency (Session 4) and an altered reliability of the visual cue (Session 5). The supplementary video shows examples of key training sessions, but note that in the experiment participants were immersed in the environment using a stereoscopic VR headset which had a wider field of view and accounted for minor head-movements, and with higher-quality sound. Another group in a control experiment also attempted to use the echo-like cues without training, but was unsuccessful, showing that the echo-like cue was indeed novel to participants. In another follow-up experiment, an additional group was trained with a novel vibrotactile cue to distance and showed similar results, demonstrating that Bayes-like integration in the main experiment was not specific to auditory or echo-like stimuli.

### Untrained Control Experiment

Performance in the untrained control experiment was worse than we would expect if participants just ignored the audio cue entirely and just pointed to the center of the response line on each trial, demonstrating that people were unable to use the echo-like audio cue without training (see SI for more details, especially Figure S1). Therefore, perception of distance via an echo-like auditory delay, although based on a natural physical relationship, was not an existing perceptual skill for naïve participants. Given that participants could not use the single audio cue without training, we omitted any further testing to see if they could also combine it without training.

### Distance Perception via the Novel Audio Cue in Trained Participants

All analyses from this point forward are based on data from just the trained participants in the main experiment. In the initial sessions, which trained them first with 2, 3 and 5-alternative forced-choice (Session 1) and then with continuous responses (Session 2), they learned the novel audio cue and were able to use it accurately to judge distance. The correlation between the logarithm of the targets and the logarithm of the responses was above 0.80 for every participant. See SI S2 for individual graphs, see Figure 2 for an example participant, and see the Supplementary Video for example trials.

**Figure 2.**
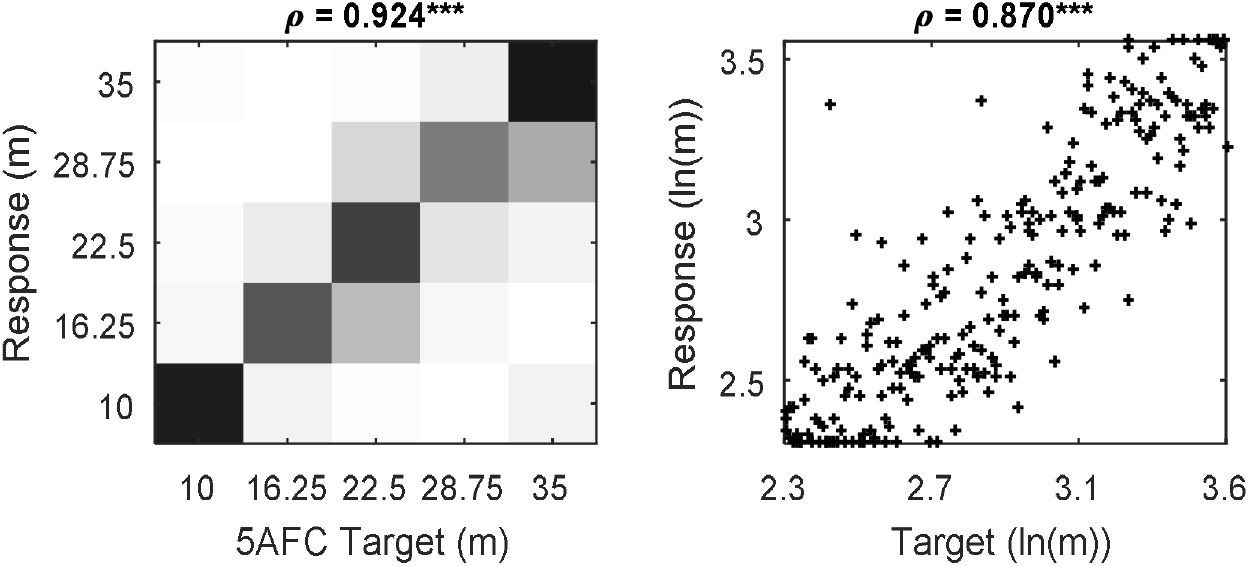
Example performance with the echo-like augmented cue. On the left are the 5AFC trials from this participant’s Session 1. Darker squares indicate more data. On the right are the continuous trials from Session 2, with each point indicating a trial. The selected participant had the median correlation between target and response in Session 2. *** p < .001.

### Main Tests for Bayes-Like Cue Combination with Vision

The two key criteria of Bayes-like cue combination were fully met by all four main tests. Exact statistics are in Table 1, and measures of variable error (the remaining error after parsing away systematic biases) and reweighting are plotted in Figures 3 and 4.

**Figure 3.**
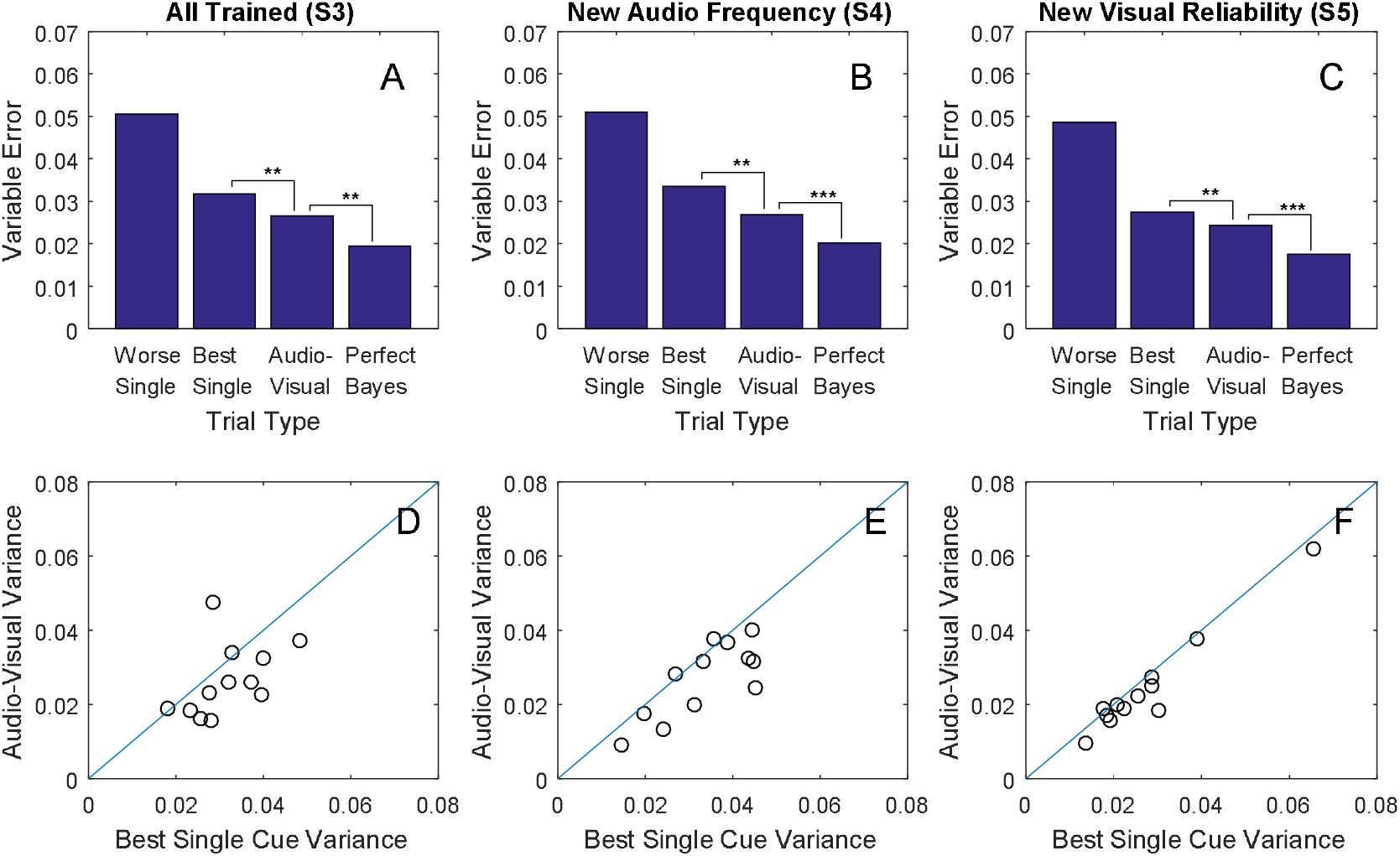
Reductions in variable error. (A-C): In each of sessions S3, S4, and S5, Audio-Visual trials (where subjects had both cues) had lower variable error than trials with the best single cue in a two-tailed sign rank test; *\*p* < .05; \*\**p* < .01; \*\*\**p* < .001. Variable Error is given on a log scale: 0.01 corresponds to a standard deviation of 10.5% of the subject-target distance, which translates to 1.1m at the near limit of 10m or 3.7m at 35m. Exact effect sizes, bootstrap confidence intervals, z-values, and p-values are in Table 1. (D-F): Each circle is an individual participant. Circles below the blue reference line indicate that variance with both cues was lower (better) than with the best single cue.

**Figure 4:**
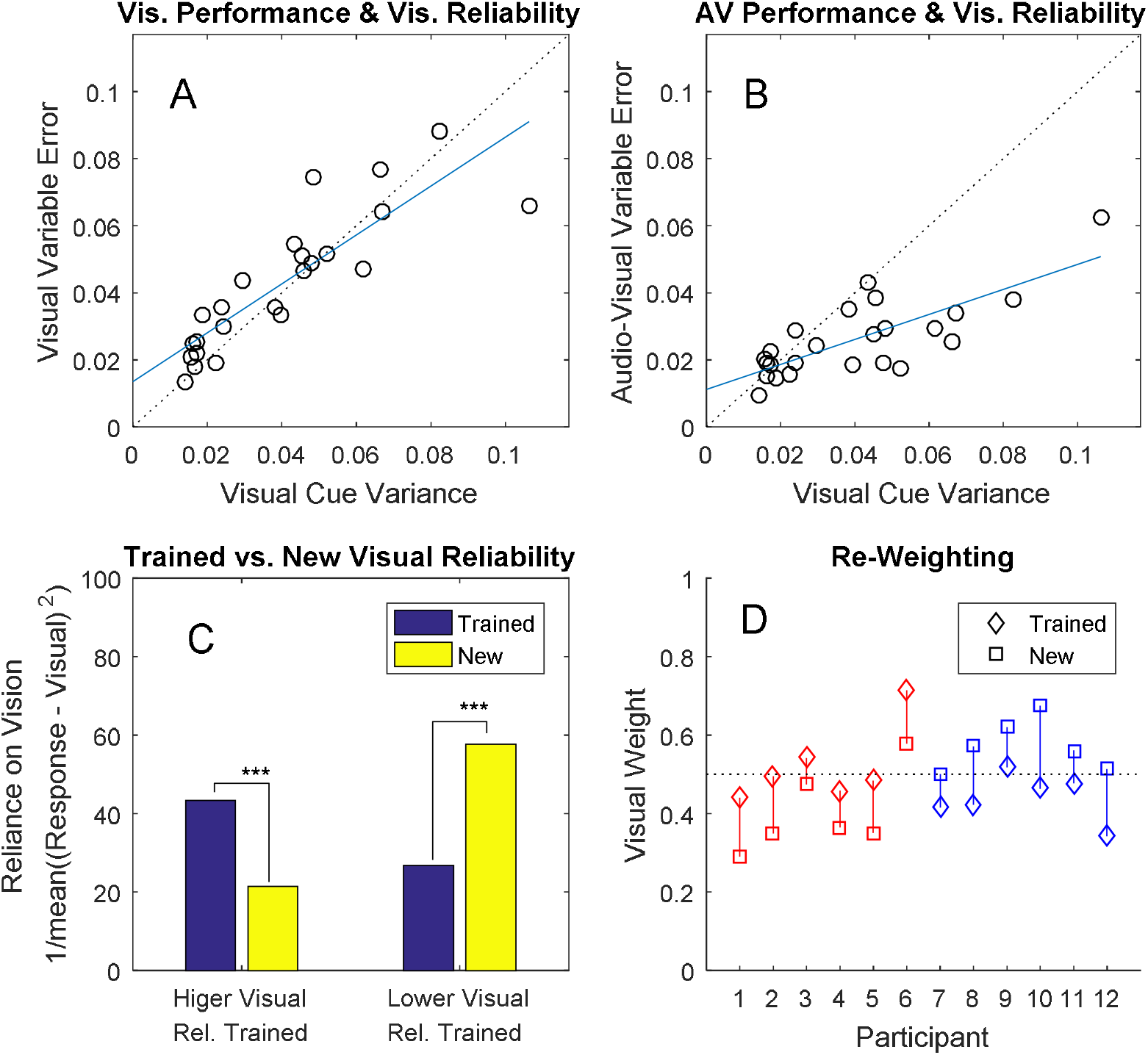
Measures of Cue Re-Weighting. (A) As a manipulation check, this shows that participants did indeed have higher variable error when the visual cue was degraded, p < .001. The blue line is the fit and the dashed line is a y = x reference line. Circles are individual participant sessions. (B) As a further manipulation check, this shows that audiovisual variable error also increased when the visual cue was degraded, p < .001. (C) On the left, participants were trained with higher visual reliability. A measure of reliance on vision (how close they pointed to the center of the visual cue) decreased when the visual reliability decreased. On the right, participants experienced an increase in visual reliability and relied on it more. (D) Inferred weights for the visual cue in Session 3 (diamonds) versus Session 5 (squares), separated into participants for whom the visual cue decreased in reliability (red) and increased in reliability (blue). Dashed reference line given at 50% weight.

**Table 1.**
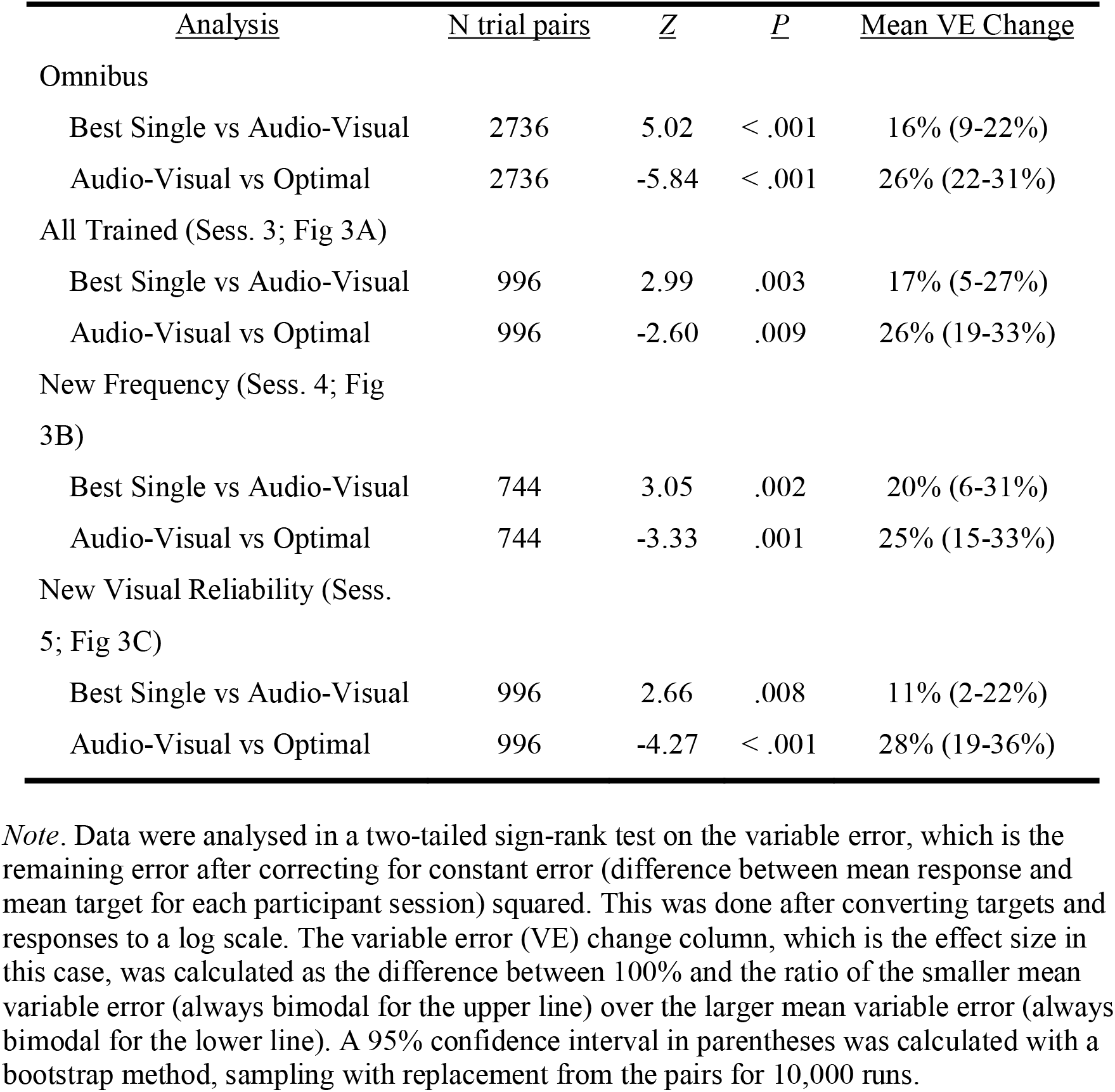
Exact statistics and Effect Sizes for Sign-Rank Tests.

### Criterion 1: reducing uncertainty (variable error) below the best single estimate when both cues are available

The first test was in Session 3, where participants were given a mix of audio-only, visual-only, and simultaneous audio-visual trials, all with feedback. A sign-rank test was favoured over a paired t-test because the distribution of variable errors is heavily skewed. The result shows that there was a significant decrease in variable error in audio-visual trials compared to the best single cue for that same participant (Table 1, Figures 3A and 3D). Thus, participants benefited from having both cues available. Specifically, they were able to make more precise judgements of distance with both cues available as compared to the best they could have done without using a cue combination strategy, thus meeting the first key criterion for Bayes-like cue combination.

A second test of this same criterion was in Session 4. Here, an untrained variation of the audio stimulus was introduced by altering the base frequency and removing feedback on all trials including this cue. The data show that even with these new, untrained audio-stimuli, and without feedback, participants still successfully lowered variable error below the best single cue on audio-visual trials in Session 4 (Table 1, Figures 3B and 3E). This independently meets the first key criterion, even in a situation where the exact stimuli have not been trained.

The third test was in Session 5. Here the reliability of the visual stimulus was altered and feedback was removed on audio-visual trials. Participants again lowered variable error for the audio-visual trials versus the best single cue (Table 1, Figures 3C and 3F). This provides a third and final independent test of the first key criterion, in a situation where participants had to adjust their audio-visual strategy based on changes to the reliability of the visual cue and therefore changes in the relative uncertainties of the two cues. This happened without feedback on the audio-visual trials.

### Criterion 2: Flexibly re-weighting cues when they change in reliability

To test for our second criterion for Bayes-like cue combination, we looked at measures of the relative weight given to the two cues across sessions. The visual cue changed from Session 3 to Session 5, increasing its variance (lengthening in the virtual world) for half of participants and decreasing its variance (shortening) for the other half. The variance of the visual cue was set to 75% (short) or 125% (long) of each participant’s observed audio variance in Session 2. On a most general level, we expected this manipulation to decrease/increase reliability of the visual cue (increase/decrease variable error). Confirming that our manipulation worked as intended, we found a significant relation between the visual cue variance and the variable error for both visual-only trials, r(22) = 0.86, p < .001, and for audio-visual trials, r(22) = 0.77, p < .001 (see Figure 4A-B), with within-subject (across-session) changes all in the expected direction. Thus, our manipulation of the visual stimulus successfully changed the reliability of the visual cue.

Under standard Bayesian theory, where participants are taking a precision-weighted average, we can predict that a cue’s weight will go down when its reliability goes down and that it will go up when its reliability goes up. The simplest way to see this is to examine how close participants responded on average to the center of the visual cue (Figure 4C). When given a higher visual variance, participants tended to point further away, and when given a lower visual variance, participants tended to point closer. A sign rank test comparing low variance (short) visual cues to high variance (long) visual cues shows a significant effect, p < .001. A more direct test for cue re-weighting was instantiated via modelling analysis as well. This model estimated the relative weights for each participant in their Sessions 3 and 5 as shown in Figure 4D. Weights for vision for all 12 changed in the expected direction, with the 99% credible interval for the mean re-weighting excluding zero. This rules out an alternative explanation whereby the correct rule is learned in a rote fashion without representing own uncertainty, as there would be zero re-weighting without further feedback (Maloney & Mamassian, 2009). It further rules out a any explanations that don’t involve averaging the two cues together, such as processing the audio cue better with the visual cue present (e.g. Thorne & Debener, 2008), since this would lead to inferred weights near zero or one.

### Additional Results

Given the approximately-Gaussian distribution of errors (S2 in SI), the optimal prediction from a Perfect Bayes integrator in Formula 1 should be a reasonable estimate of the best possible benefit from having both cues available (Ernst & Banks, 2002):

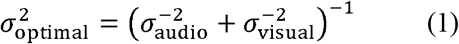

Participants fell short of this prediction in every session and regardless of which cue was more reliable (Figure 3A-C and Table 1). This suggests that while Bayes-like cue combination was occurring, it was not yet tuned optimally.

SI provides additional analyses that do not change the main interpretation, but suggest that the main test results are robust to reasonable variations in exactly what is entered into the analysis: total error (squared distance between target and response) and average variable error for each participant.

### Vibrotactile Experiment

In the SI we also present an additional dataset where the study here was repeated with a novel vibrotactile cue to distance, omitting Session 4 (since there was no audio frequency to alter and the vibrotactile device did not have a similar option available). In these data, with the same analytic methods (see S4 especially), both criteria for Bayes-like combination were also met. This shows that rapid cue-combination via newly learned cues is not only specific to auditory or echo-like signals, or those based on natural physical relationships.

## Discussion

Both key criteria for Bayes-like cue combination were fully met by all analyses. Both cues must have been used when they were both available, because otherwise we would not expect systematic improvements in precision when compared to the best single cue. Participants must have had some representation of their own uncertainty, because otherwise they would not have been able to re-weight the cues when their relative reliability changed and feedback was removed. Participants must have averaged the two cues, rather than have the presentation of one cue enhance the processing of the other cue, because otherwise the enhancing cue would receive no inferred weight. This was true across the four main planned analyses in the Results and also across several variations in SI – including a replication using a novel vibrotactile cue instead of the auditory echo-like cue.

These results are a proof-of-principle that learning to combine a new sensory skill with a familiar sensory skill is well within the realm of human possibility. This point is crucial because it means that new sensory skills need not replace familiar sensory skills.

Humans whose senses are augmented stand to gain the Bayesian benefits of incorporating the new and standard sensory information, rather than having to choose between them. This discovery suggests that techniques like echolocation (Kolarik et al., 2014; Stroffregen & Pittenger, 1995; Thaler & Goodale, 2016) and devices translating information from one modality into another (Abboud et al., 2014; Maidenbaum et al., 2014; Meijer, 1992) might not only hold promise for people with complete sensory loss (e.g. total blindness), but could in principle be a useful aid for less severe levels of impairment – for example, moderate vision impairments, estimated to affect 214 million people alive today (Vos et al., 2016). Our results also suggest an optimistic outlook on sensory augmentation more broadly, allowing people to use novel signals efficiently during specialist tasks – for example, if novel auditory cues to position during brain surgery (Woerdeman, Willems, Noordmans, & van der Sprenkel, 2009) were combined with visual information.

This result is unexpected under previous models of multisensory learning (Daee et al., 2014; Weisswange et al., 2011), and it follows that these models will need to be updated to account for these findings. This result is further surprising given that the most comparable previous study, with a new sense of turning, found cue alternation (choosing between cues trial to trial) rather than cue combination (Goeke, Planera, Finger, & König, 2016). By contrast, we find evidence for Bayes-like combination, both in the main experiment using an echo-like cue, and in the additional experiment (see SI, especially S4) showing the same results with a vibrotactile cue. The difference in results might be due to certain choices in the methods. Regardless, the present study provides a basis for future studies teaching sensory augmentation skills to moderate sensory loss patients, and examining if benefits can move from lab to everyday life.

However, our findings also come with a notable limitation. Our analyses suggest that our participants achieved less than the optimal variance reduction predicted for a perfect Bayesian integrator (Figure 3A-C; 4B-C; Table 1). In contrast, studies with highly-familiar cues often show performance indistinguishable from optimal (Alais & Burr, 2004; Ernst & Banks, 2002; Hillis et al., 2004; Knill & Saunders, 2003); though see also (Rahnev & Denison, 2017). One potential explanation is that perfect Bayesian integrators must choose cue weights exactly proportionate to cue reliabilities, but our participants may not have yet learned to judge cue reliabilities precisely. It is unknown how rapidly and accurately humans learn reliabilities of novel cues. Related studies on the acquisition of novel perceptual priors show that reliabilities are not always learned accurately in limited training sessions – for example, in 1200 trials (Bejjanki et al., 2016), similar to here. Another potential explanation is that cue combination depends on inferring that two cues share a common cause (Shams & Beierholm, 2010). This can limit combination with cues that are perceived as discrepant, and with unfamiliar cues in particular, this might affect combination. We predict, but cannot empirically confirm at this time, that near-optimal combination would be found after much more extensive training.

The present study, conducted entirely with adults, may counter-intuitively be one of the best ways to answer a developmental question as well. Adults generally combine a variety of independent cues that children under 10-12 years do not, such as visual texture and disparity cues to surface slant (Nardini et al., 2010). This brings up a question about mechanisms; when comparing a 20-year-old to an 8-year-old, there are differences in both general maturation (e.g. of neural substrates) and in how much cue-specific experience they have had (e.g. how many slanted surfaces they have looked at). To what degree is each of these responsible for the developmental difference in cue combination behaviour? Current models place the emphasis entirely on cue-specific experience (Daee et al., 2014; Weisswange et al., 2011) and neglect maturational changes. To examine this empirically, the two factors need to be dissociated. It would not be ethical to deprive children of cue-specific experience while they matured, and likely not feasible to meaningfully increase their rate of cue-specific experience for ubiquitous properties such as surface slant. The current study instead created a situation where people with full maturation (adults) had no cue-specific experience (new sensory skill). Cue combination was learned rapidly, emphasizing the importance of maturation as a key difference that allows cue combination behaviour to be expressed. There are proposals, but as yet no clear consensus, on how reliability-weighted cue combination is implemented even in the adult brain (Carandini & Heeger, 2012; Ohshiro, Angelaki, & DeAngelis, 2011; Rosenberg, Patterson, & Angelaki, 2015). Better understanding these mechanisms and how they mature to support flexible cue combination abilities is a crucial question for further research.

In conclusion, Bayes-like cue combination can extend to augmented or substituted senses. This suggests an optimistic outlook on augmenting the human senses, with potential applications not only to overcoming sensory loss, but also to providing people with completely new kinds of useful signals. To make this even more exciting, it can also happen rapidly. Further study will be needed to examine the full extent of this ability and the circumstances under which it is shown.

## Materials and Methods

### Participants

For the main study, 12 healthy adult participants were recruited using the Durham Psychology Participant Pool and posters around Durham University. There were 2 males and a mean age of 24.9 years (age range 19.8-39.8 years; standard deviation 5.5 years). A single pilot participant was also run, whose data were not included in the main analysis but instead used for a power analysis. That analysis found over 95% power for the primary tests (Table 1) with 12 participants with the same level of noise and bias in their estimates. We also ran a control experiment to determine if the echolocation cue was indeed new (i.e. not useful without training). For the control experiment, an additional 12 participants were recruited, including 1 male and with a mean age of 19.4 years (age range 18-20 years; standard deviation 0.66 years). Power calculations suggested that a difference between control and training should be significant >95% of the time if the control group was not using the audio cue at all. Participants were paid £8 per hour in both experiments. Ethics were approved by the Durham University Psychology Department’s Ethics Committee and informed consent was obtained in writing. Methods were carried out in accordance with their guidelines and relevant regulations. Participant and other details for the additional vibro-tactile experiment are found in the SI.

### Apparatus

#### Virtual environment

A custom seascape was created in WorldViz Vizard 5 (Santa Barbara, CA, USA) and presented using an Oculus Rift headset (Menlo Park, CA, USA).

This seascape contained a large flat blue sea, a ‘pirate ship’ with masts and other items, a virtual chair, and a friendly cartoon whale introduced with the name “Patchy” (see Figure 1). Participants were seated 4.25 m above the sea. The response line (range of possible positions for the whale) stretched out from the bow of the ship, and was marked by periodic variations in the color of the sea. A pair of black bars marked the 10 m and 35 m points, which were the nearest and furthest possible responses. Distances along the line could be judged visually via perspective and height-in plane (see Figure 1A, 1C) as well as, in theory, stereo disparity (although stereo information at the distances used is of limited use). Patchy gave written instructions via a white speech bubble (example, Figure 1A). To respond, participants set the position of a 3D arrow that touched the sea surface. The sea surface remained still and the ship did not move. See the Supplementary Video for example trials.

#### Headset

The Oculus Rift headset has a refresh rate of 90 Hz, a resolution of 1080 x 1200 for each eye, and a diagonal field of view of 110 degrees. Participants were encouraged to sit still and look straight ahead during trials but did not have their head position fixed. The Rift’s tracking camera and internal accelerometer and gyroscope accounted for any head movements in order to render an immersive experience.

#### Audio equipment

Sound was generated and played using a MATLAB program with a bit depth of 24 and a sampling rate of 96 kHz. A USB sound card (Creative SoundBlaster SB1240; Singapore) was attached to a pair of AKG K271 MkII headphones (Vienna, Austria) with an impedance of 55 ohms.

#### Controller

Participants used an Xbox One controller (Redmond, WA, USA). They only used the left joystick and the A button. Pressing the other buttons did not have any effect on the experiment.

### Stimuli

#### Audio cue

The audio stimuli were created by first generating a 5ms sine wave either 4000Hz or 2000Hz in frequency with an amplitude of 1. Half of participants experienced the higher frequency in Sessions 1-3 and 5, switching to the lower in Session 4, and vice-versa (see below). The first half-period of the wave was scaled down by a factor of 0.6. An exponential decay mask was created starting after 1.5 periods and ending at 5ms. The exponent was interpolated linearly between 0 and −10 over that period. This was all embedded in 1s of silence, with a 50ms delay before the sound appeared. An exact copy of the sound was added after an appropriate delay, calculating the distance to the target divided by the speed of sound (approximated at 350m/s), then times two (for the emission to go out, and also to come back). With a minimum distance of 10m, the two sounds (clicks) never overlapped (although it is possible that subjects experienced them as one sound). Real echoes contain more complex information, including reductions in amplitude with distance, but we chose to make delay the only relevant cue so that we could be certain which information all participants were using. Our stimuli also allowed us to use range of distances at which real echoes are typically very faint, minimizing the scope for participants to have prior experience with them.

#### Visual cue

Vision provides a wide variety of depth cues that interact heavily (Howard & Rogers, 2008) and require various techniques to isolate fully. However, we wanted to frame the task overall as a reasonably naturalistic ‘hide and seek’ game with a social agent. This is in conflict with these isolation techniques. We also wanted to have precise control over how useful this cue was so that we could match it well to each participant’s learning with the echo-like cue. To achieve this, the visual cue had strong external noise and negligible internal noise.

The visual cue was a fully 3D array of 256 ‘bubbles’ (translucent white spheres with a radius of 0.15 m and 50% opacity) arranged to show a mirrored log-normal distribution perpendicular from the line on which the whale appeared (Figure 1C). This was generated by first lining the outer edges with the bubbles and then filling in the interior area such that no two bubbles touched each other. The result looks like a violin plot of a log-normal distribution. This was arranged such that the target was always an actual draw from the distribution on display, i.e. the whale’s true hiding place was under one of the bubbles. This means that when given only visual information, the error-minimising (ideal observer) strategy is to aim at the center of the bubble distribution where it is most dense (Figure 1C). Given additional positional information via an independent second (e.g. auditory) cue, the ideal observer strategy is a reliability-weighted average of the center of the bubble distribution and the position indicated by the other cue. The variance of the visual cue (i.e. the ‘length’ of the bubble distribution along the response line) was calculated for each participant based on their auditory-cue variance, and changed in the final session; see below.

### Procedure for the Training Condition

Participants were told that that they were going to play a kind of ‘hide and seek’ game with Patchy the whale. On each trial, they received the visual and/or auditory cues and then used the joystick to move an arrow to try to point as close as they could to where Patchy was hiding. There was no time limit. Participants were not given any additional information while responding and were not allowed to hear the audio stimulus again. The visual stimulus remained static on the sea while they were responding. For trials with feedback, Patchy appeared at his true location with a speech bubble indicating the error as a percentage of his true distance away from the ship (e.g. if he was 10 m away and they pointed 12 m away, they would see +20%). The text was provided in addition to seeing the whale in case there was any confusion about which point on the whale, from its head to its tail, was the precise point that participants were attempting to localize. (It was actually in the whale’s center.)

#### Session structure

The full script of the different sessions is detailed in the SI and briefly explained here. Example trials can also be seen in the Supplementary Video. Session 1 and 2 taught the audio cue to the participants. Session 1 (300 trials) started with 50 two-alternative-forced choice (2AFC) task trials, choosing among a point at the nearest vs farthest limit of the line. It then progressed to 100 trials of 3AFC (including the midpoint) and 150 trials of 5AFC (including the 25% and 75% points). The correct targets were as evenly distributed as possible. The whale surfaced after every trial to give accurate feedback.

Session 2 (300 trials), and all further sessions, began with a short 40-trial warm-up of 2-, 3-, and 5-alternative-choice trials to remind participants how the audio cue works. In the second session, this proceeded to 250 audio-only trials with feedback, now with a continuous response (all positions along the line), with the targets spaced evenly on a log scale. This staging of training, from 2AFC to continuous, was done to help scaffold the participants towards consistent performance. Common to Sessions 2-4, the last 10 trials were used to introduce what would happen in the next session.

Session 3 (299 trials; 83 matched triplets) tested for cue combination by showing participants the visual cue only, the audio cue only, or both together. Feedback was given throughout. This was used to test the first criterion for Bayes-like combination, whether the variance of combined estimates was lower than the variance of single-cue estimates.

Session 4 (298 trials; 62 matched triplets) tested for the ability to generalize the echo-like cue to a new emission, specifically a click with a different frequency of the amplitude modulated sine wave. The session was very similar to Session 3 except that all trials with the new sound were not given feedback. This further tested the first criterion. This manipulation was inspired by the finding that actual echolocation users have some meaningful variation in exactly what frequencies they emit from click to click (Zhang et al., 2017).

Session 5 (300 trials; 83 matched triplets) tested for the ability to generalize to a new reliability level of the visual cue. It was much like Session 3 except that the log-normal distribution displayed by the visual cue changed in variance. Feedback was given on the visual-only trials to allow participants to see how the reliability had changed, but not on the bimodal trials, so that they had to infer the correct weights from their knowledge of each cue’s reliability. This was used to test the second criterion, reliability-based reweighting of cues.

#### Stimulus matching for sessions 3 through 5

Each of the last three sessions involved audio-only, visual-only, and audio-visual trials. These were generated in triplets, starting with the audio-visual trials. The centers of the visual distributions were spaced evenly on a log scale and the true target was drawn from the distribution they showed. The audio cue was generated to signal the target exactly. The standard deviation of the visual cue was either 75% or 125% of the error in audio-only responses during Session 2. The 75% and 125% figures were chosen because they are close enough to 100% that we still expect a substantial cue combination effect (if one is vastly more reliable than the other, only a vanishingly small effect is expected), but far enough from each other that we expected to be able to measure changes in cue weighting. Half of participants were shown visual cues with the 75% standard deviation for Session 3 and 4, then 125% for Session 5, and the other half vice versa. A matching audio-only trial was made by just removing the visual cue, and a visual-only by just removing the audio cue. This means that when we compare the single-cue trials to the dual-cue trials within each triplet, there was nothing different except the other unused cue in the single-cue trials. Order of presentation was randomized.

### Procedure for the Untrained Control Experiment

These data were collected to be sure that the echolocation-like sensory skill was genuinely new and not familiar from everyday experience with reflected sound. Participants were told that we wanted to know if people without echolocation training had any intuitions about how echolocation might work, that the sounds they were going to hear were echoes that trained participants could use to estimate distance, and that their goal was to respond as close to Patchy as possible despite never seeing where he actually was. The stimuli were exactly the same as the continuous-response section of Session 2 (250 trials, audio only, distributed evenly on a log scale), but there was no feedback given in any part of this experiment. There was only one session.

### Data Processing

The data from the second half of Session 1 (which was just training with the audio cue) and the continuous trials of Session 2 (further audio cue training) were transformed onto a log-scale to account for the way that auditory delays become harder to perceive in linear terms as the delays become longer (Getty, 1975).

For sessions 3-5, targets and responses were again transformed onto a log-scale for the same reason. For the first three main tests (out of four), the log error was parsed into constant error (the bias component) and variable error (the precision component). The constant error was calculated separately for each participant, session, and trial type combination (12 participants x 3 post-training sessions x 3 trial types = 108 bias corrections). The specific formula was:

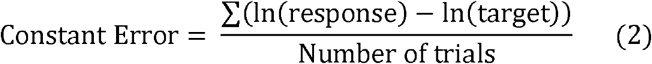

The variable error was calculated for each trial for the main analysis. (An additional analysis is presented in SI, using an average variable error instead.) The specific formulas was:

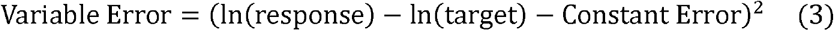

The best single cue was determined at the level of participant and session, averaging the variable errors over all targets within each modality and selecting the lower of the two. The variable error for each bimodal trial was paired for analysis with the variable error from the best single cue trial for that same target, same participant, and same session. This gives us the most power to detect possible changes in variable error with the addition of the worse cue (which is the only change across these pairs), and thus the best test of the first criterion.

Simulations were also performed to be sure that this method does not inflate the type I error rate, suggesting that, if anything, the analysis method here is slightly conservative against finding differences in variable error (roughly 4.8% of p-values below the nominal .05 threshold). Statistics calculations were performed in Matlab 2016a by MathWorks.

### Re-Weighting Model

This model was used to estimate weights and to assess their pattern of change from Session 3 to Session 5 when the visual cue reliabilities changed. The model has two layers. The lower layer was designed to let us estimate the weight given to each cue within each participant and session. The central equation here was:

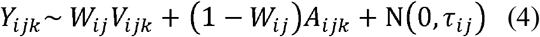

where *i* gives the participant, *j* gives the session, *k* gives the trial number, *Y* is the response, *W* is the weight given to the visual cue, *A* is the placement of the audio cue, *V* is the placement of the visual cue, and τ is the precision of responses. In other words, participants respond at a weighted average of the two cues plus some noise, with the weights as free parameters.

The higher layer of the model was designed to relate each person’s cue weights in Session 3 to their cue weights in Session 5, specifically to see if it fits a pattern that matches Bayesian predictions (higher visual reliability leads to higher visual weight). The central equations here were:

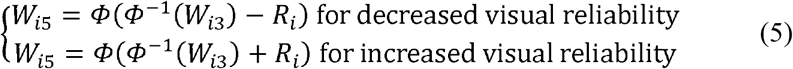

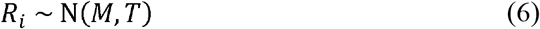

where *R* is the reweighting effect, *M* is the mean reweighting, and *T* is the precision of reweighting effects. In other words, a positive value for *M* fits Bayesian predictions, but a value of zero for *M* fits the learning of a static weighting.

The priors for each parameter were:

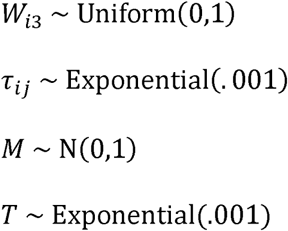

These were fit in WinBUGS (Lunn, Thomas, Best, & Spiegelhalter, 2000) with 6 independent chains consisting of 25,000 used samples and 5,000 burnin-in samples each (total 150,000 used samples). In the WinBUGS implementation, an exponential distribution is given its rate parameter, so the prior on *T* had a mean of 1,000 (extremely vague).

Note then that several possible assumptions are not ‘built-in’ to the model. First, it is capable of fitting a value of W arbitrarily close to 0 or 1. At W=0 or W=1, the model simplifies to a cue selection or weak interaction model. Second, there is no requirement that it fits a specific audio-visual variance, so it is free to fit a wider range of weights and variances than just the ones predicted by standard Bayesian reasoning. Third, the model is perfectly capable of finding a mean weight change *M* that is very near zero and includes it in its credible interval. To check this, we purposefully switched participants 4-6 with participants 10-12, which fit *M* with an interval including zero. Fourth, there is no hierarchical structure to the weights in Session 3, so *M* is only capturing mean changes rather than the mean of weights in Session 5.

## Materials and Data Availability

The data are attached as SI. The code is available on request to the first author.

## Acknowledgements

We would like to acknowledge support through grant ES/N01846X/1 from the UK Economic and Social Research Council. Thanks to Hannah Roome, Alysha Chelliah and Claudia Bowman for help with piloting.

## Author Contributions

JN, LT and MN conceived of the project. LT provided the audio stimuli. JN programmed the rest of the experiment. LW collected the data. JN analysed the data and drafted the manuscript. LT and MN provided comments. MN also held a supervisory role in the project.

## Competing Interests

The authors declare that they have no competing interests.

